# Synergizing Bayesian and Heuristic Approaches: D-BPP Uncovers Ghost Introgression in *Panthera* and *Thuja*

**DOI:** 10.1101/2025.06.27.662067

**Authors:** Yang Yang, Xiao-Xu Pang, Ya-Mei Ding, Bo-Wen Zhang, Wei-Ning Bai, Da-Yong Zhang

## Abstract

Hybridization involving extinct or unsampled ("ghost") lineages profoundly influences species’ evolutionary histories, but detecting such introgression remains methodologically challenging. We introduce D-BPP, a novel framework that integrates the heuristic *D*-statistic (or ABBA-BABA test) with Bayesian phylogenomic inference (implemented in BPP) to efficiently infer phylogenetic networks. In D-BPP, we first employ the *D*-statistic to rapidly identify candidate introgression events on a predefined binary species tree; then we leverage the Bayesian test in BPP to rigorously validate these candidates and sequentially add them to the species tree, retaining only those events with strong statistical support. If the species tree is ambiguous, D-BPP identifies the most probable tree by evaluating competing topologies through Bayesian model comparison of their corresponding introgression models. Critically, our framework excels at detecting ghost introgression, which is often unidentifiable or overlooked by existing methods— whether heuristic or full-likelihood. Applied to genomic datasets from *Panthera* (big cats) and *Thuja* (conifers), D-BPP uncovered previously undetected ghost introgression events in both clades, underscoring the pervasive role ghost lineages have played across diverse taxa. By uniquely combining the computational efficiency of heuristic *D*-statistics with the robust statistical rigor of full-likelihood Bayesian inference, D-BPP deciphers complex hybridization patterns obscured by conventional methods, providing a powerful tool for accurately reconstructing phylogenetic networks.

## Introduction

Speciation, extinction, and hybridization are fundamental processes that together shape the evolutionary trajectories of plants and animals. These processes often interact in complex ways that violate the assumptions of strictly bifurcating phylogenies, necessitating the adoption of phylogenetic networks as a more realistic framework for depicting species relationships (Yu et al. 2014; Degnan 2018; Blair and Ané 2020; Solís-Lemus and Tiley 2025). Unlike traditional trees, networks explicitly accommodate gene flow not only among extant lineages but also from ghost lineages—extinct or unsampled taxa whose introgression leaves detectable genomic signatures (Ottenburghs 2020; Tricou et al. 2022a, b; Pang and Zhang 2023; Tiley et al. 2023; Pang and Zhang 2024). A growing body of evidence suggests that such ghost introgression is widespread across the tree of life, with recent examples reported in diverse taxa (Pawar et al. 2023; Zhang et al. 2024; Baraf et al. 2025; Shen et al. 2025). Nonetheless, detecting these events remains a major methodological challenge (Pang and Zhang 2024).

A wide range of methods have been developed to directly infer phylogenetic networks from genomic data, encompassing both heuristic and full-likelihood strategies as reviewed in Jiao et al. (2021) and Hibbins and Hahn (2022). Full-likelihood approaches, which make use of multi-locus sequence alignments, offer superior statistical power for detecting gene flow (Yang and Flouri 2022; Pang and Zhang 2024). However, their application to large datasets is hindered by substantial computational demands (Wen and Nakhleh 2018; Zhang et al. 2018). Meanwhile, heuristic approaches based on summary statistics are generally limited to simple level-1 networks (i.e., those with non-overlapping reticulations), and more complex networks may be statistically non-identifiable (Yu and Nakhleh 2015; Solís-Lemus and Ané 2016; Allman et al. 2019; Kolbow et al. 2025; Kong et al. 2025). To address these issues—poor scalability and network oversimplification/non-identification— researchers often first estimate a bifurcating species tree and then add reticulations to it using tree-guided introgression detection methods such as the *D*-statistic (ABBA-BABA test) (Hibbins and Hahn 2022). However, this strategy faces two main challenges. The first concerns species tree estimation. Gene flow, particularly between non-sister lineages, can distort phylogenetic signals and bias species tree inference (Mallet et al. 2016; Jiao et al. 2020; Pang and Zhang 2023; Dinh and Baños 2025). Under such conditions, accurate inference of speciation order becomes difficult, and errors in the species tree can lead to incorrect networks. A potential solution, nonetheless, may exist: following Santos et al. (2025) among many others, one may evaluate multiple introgression models constructed from alternative species-tree topologies through Bayesian model comparison to identify the most plausible phylogeny.

The second challenge concerns the detection of introgression events. Among many heuristic methods, the *D*-statistic (Green et al. 2010; Durand et al. 2011) has remained the most widely used since its introduction (Dagilis et al. 2022), which leverages genome-wide site pattern counts in a species quartet (three ingroup taxa and one outgroup) with a specie tree (((P1,P2),P3),O). Denoting the ancestral allele as “A” and the derived allele as “B”, there are two discordant site patterns: ABBA and BABA. Under the null hypothesis of no gene flow, both ABBA and BABA patterns can only be attributed to incomplete lineage sorting (ILS) and thus should be observed in equal numbers. Therefore, significant deviations from this expectation, resulting in the statistic 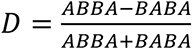 significantly different from zero, provides convincing evidence for the presence of gene flow, usually interpreted as introgression between two of the three ingroup lineages. The key strengths of this method lie in its ability to provide a formal statistical framework for discriminating phylogenetic incongruence caused by ILS from that caused by gene flow (Zheng and Janke 2018; Kong and Kubatko 2021), as well as its high computational efficiency in handling genome-scale datasets. However, simulations by Tricou et al. (2022b) revealed that introgression from unsampled outgroups can also generate significant *D*-statistic signals, making the usual interpretation of a significant *D*-statistic unreliable.

By contrast, full-likelihood methods can distinguish among alternative introgression scenarios that may produce similar *D*-statistic signals (Pang and Zhang 2024), but their use has been limited by computational demands. Ji et al. (2023) recently addressed this issue by developing a tree-guided Bayesian test for introgression within the BPP program (Flouri et al. 2020), which employs the Savage–Dickey density ratio (Dickey 1971) to efficiently compute Bayes factors when the two compared models are nested. This approach avoids computational bottlenecks associated with traditional full-likelihood methods, such as cross-model searches and marginal likelihood calculations (Huang et al. 2022; Ji et al. 2023). The method has since been applied in multiple studies (Ji et al. 2023; Ji et al. 2025; Zhu et al. 2025) to infer reticulate evolution by systematically evaluating predefined ingroup introgression events and retaining those with strong Bayes factor support. However, these analyses assume introgression occurs only among ingroup members, neglecting gene flow from ghost lineages—extinct or unsampled taxa that introgress into extant ingroup members—despite their potential importance in shaping the evolutionary histories of clades like *Panthera* (big cats) and *Thuja* (arborvitae).

To overcome these challenges, we developed D-BPP—a unified framework combining the computational efficiency of the *D*-statistic with the statistical efficiency of the Bayesian test implemented in BPP—to jointly resolve species divergence histories and introgression patterns in reticulate systems. We applied D- BPP to two empirical datasets: *Panthera* (Santos et al. 2025) and *Thuja* (Li et al. 2022), analyzing each group under three competing species-tree topologies. Our analyses clarified their species relationships and revealed complex reticulate histories involving multiple introgression events from both extant and ghost lineages, most of which were either overlooked or unidentifiable in previous studies.

## Results

### D-BPP: A Unified Bayesian–Heuristic Framework for Phylogenetic Network Inference

To resolve complex evolutionary relationships shaped by introgression, we developed D-BPP—a modular, four-step framework that integrates heuristic and Bayesian methods (Fig. 1). The model is reconstructed under the multispecies coalescent with introgression (MSci) framework (Meng and Kubatko 2009; Flouri et al. 2020), which accommodates both incomplete lineage sorting (ILS) and introgression.

**Fig. 1.**
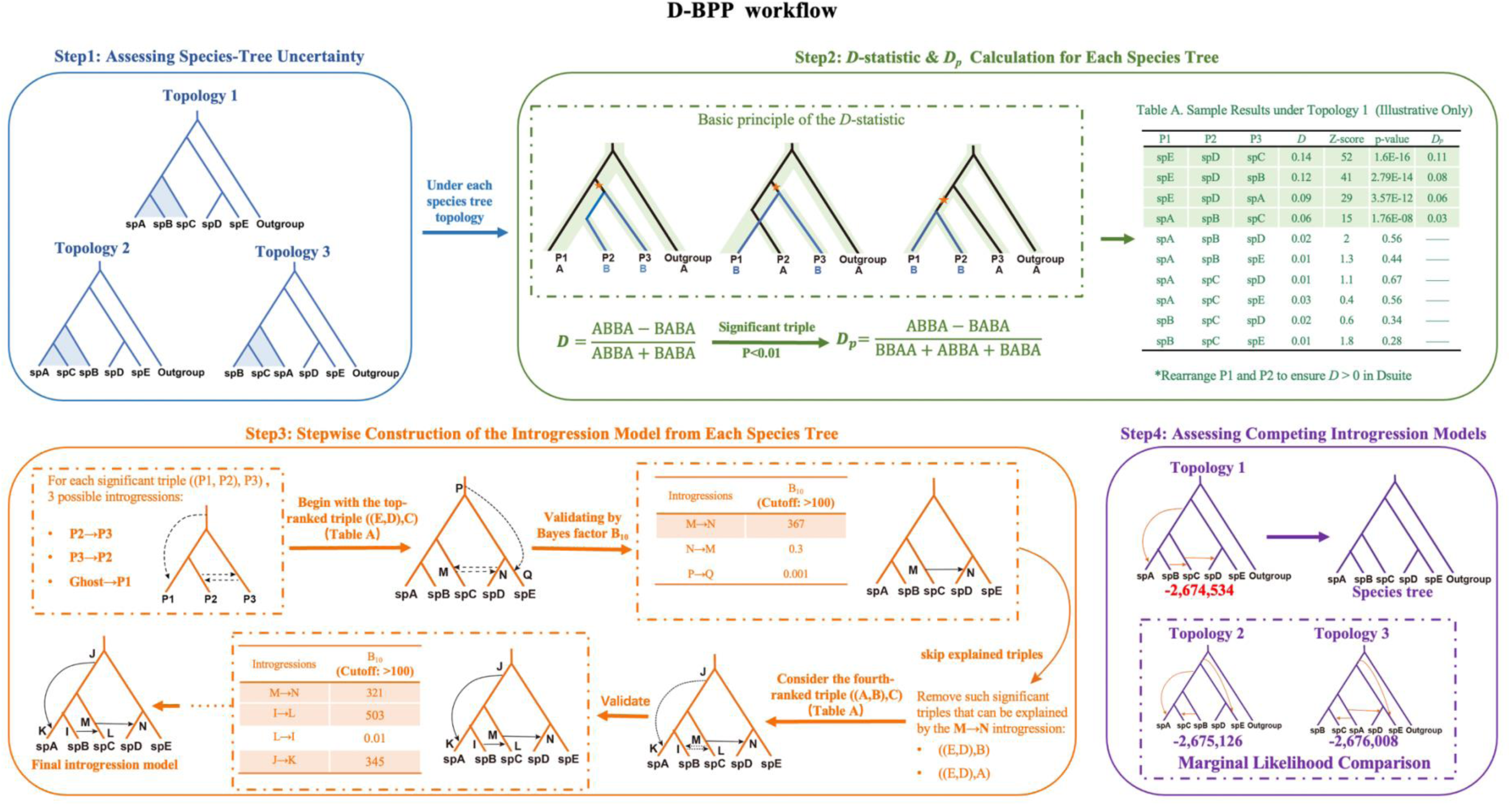
Overview of the D-BPP workflow for stepwise construction of introgression models. T D-BPP integrates heuristic D-statistic screening and full-likelihood Bayesian inference into a unified four-step workflow. Step 1: Assessing Species-Tree Uncertainty (blue section)—In a representative case involving five ingroup taxa and one outgroup, species D and E are assumed to form a well-supported sister clade. Under this constraint, there are three possible species-tree topologies for species A, B, and C, each representing an alternative hypothesis regarding their order of divergence. These topologies serve as competing backbone trees for subsequent introgression modeling. Step 2: *D*-statistic & *D_p_* Calculation for Each Species Tree (green section) — For each specified species-tree topology, all possible taxon triples ((P1, P2), P3) are generated. Within each triple, the *D*-statistic is applied to determine the presence of introgression. The mutation giving rise to the derived allele, denoted as “B,” is marked with an orange star. Significant triples (p-value < 0.01) are retained, and a normalized *D_p_* is calculated to prioritize triples with the strongest introgression signals. Table A presents illustrative results under Topology 1, where top-ranked triples include ((E,D),C), ((E,D),B), ((E,D),A), and ((A,B),C). Step 3: Stepwise Construction of the Introgression Model for each Species Tree (orange section)—For each significant triple ((P1, P2), P3), three possible introgression events are tested in BPP: (1) introgression from P2 to P3, (2) introgression from P3 to P2, and (3) ghost introgression to P1. Taking Topology 1 as an example, analysis begins with the highest-ranked triple ((E,D),C), which yields three candidate events: introgression from species C to D (M→N), from D to C (N→M), and ghost introgression to species E (P→Q). Nodes M, N, P, and Q denote specific nodes where introgression is hypothesized to occur. Each introgression event is evaluated using Bayes factors (B₁₀ > 100 indicates decisive support), and the well-supported events are incorporated sequentially into the growing network model. For instance, the M→N event received decisive support and was retained. Remaining triples whose signals are explained by previously incorporated events (e.g., ((E,D),B) and ((E,D),A)) are excluded. The next unexplained triple, ((A,B),C), is then evaluated following the same procedure. This iterative process continues until all significant triples are well explained. The final introgression model under Topology 1 included three events. The same procedure was applied to Topologies 2 and 3 to construct their respective introgression models. Step 4: Assessing Competing Introgression Models (purple section) —The marginal likelihoods of the final models inferred under each species-tree topology were compared to identify the best-supported introgression events. The species tree associated with the highest marginal likelihood was considered the most plausible representation of the true species divergence history.

In Step 1, we assess species-tree uncertainty by identifying as many plausible topologies as we can. Instead of relying on a single fixed species tree—as is often done (e.g., (Ji et al. 2023; Ji et al. 2025))—we examine multiple plausible topologies, acknowledging that both ILS and gene flow can obscure phylogenetic signals. In practice, conflicting phylogenies may arise from different genetic markers (e.g., mitochondrial vs. nuclear DNA) and/or analytical methods (e.g., coalescent versus concatenation). With whole-genome data, it is now common to reconstruct sliding- window phylogenies along the genome (e.g., (Santos et al. 2025)), often yielding multiple plausible topologies that require further evaluation.

In Step 2, for each candidate species-tree topology, we apply the *D*-statistic (Green et al. 2010; Durand et al. 2011) to all valid species triples in the form ((P1, P2), P3), using an outgroup to polarize derived allele states. Under the null hypothesis of no gene flow, the two discordant site patterns, ‘ABBA’ and ‘BABA’, are expected to occur at equal frequencies. In Dsuite (Malinsky et al. 2021), P1 and P2 are ordered such that the *D*-statistic is always positive. A non-zero *D*-statistic, typically resulting from a significant excess of ABBA over BABA, is commonly interpreted as gene flow between P2 and P3—either from P3 to P2 (inflow, Fig. 2A) or vice versa (outflow, Fig. 2B). However, Tricou et al. (Tricou et al. 2022b) demonstrated that introgression into P1 from an unsampled “midgroup” lineage—situated between the ingroup and outgroup—can also produce a significant *D*-value (Fig. 2C). We further demonstrated that ghost lineages diverging before the outgroup (“basal” ghost lineages, Fig. 2D) or originating from the outgroup itself (“outgroup-derived” ghost lineages, Fig. 2E) can likewise yield significant *D*-statistic (see Supplementary Text for a mathematical proof). Thus, each significant *D*-statistic may reflect one of three candidate introgression events, i.e., inflow, outflow, and ghost introgression (involving three distinct relative positions of the ghost lineage with respect to the outgroup; Fig. 2). For each significant triple, we computed 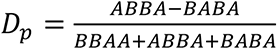 (Hamlin et al. 2020), a statistic conceptually more aligned with genome-wide introgression proportion (Hibbins and Hahn 2022). Significant triples were ranked in descending order of their *D*_p_ values (Fig. 1), facilitating efficient prioritization for downstream testing.

**Fig. 2.**
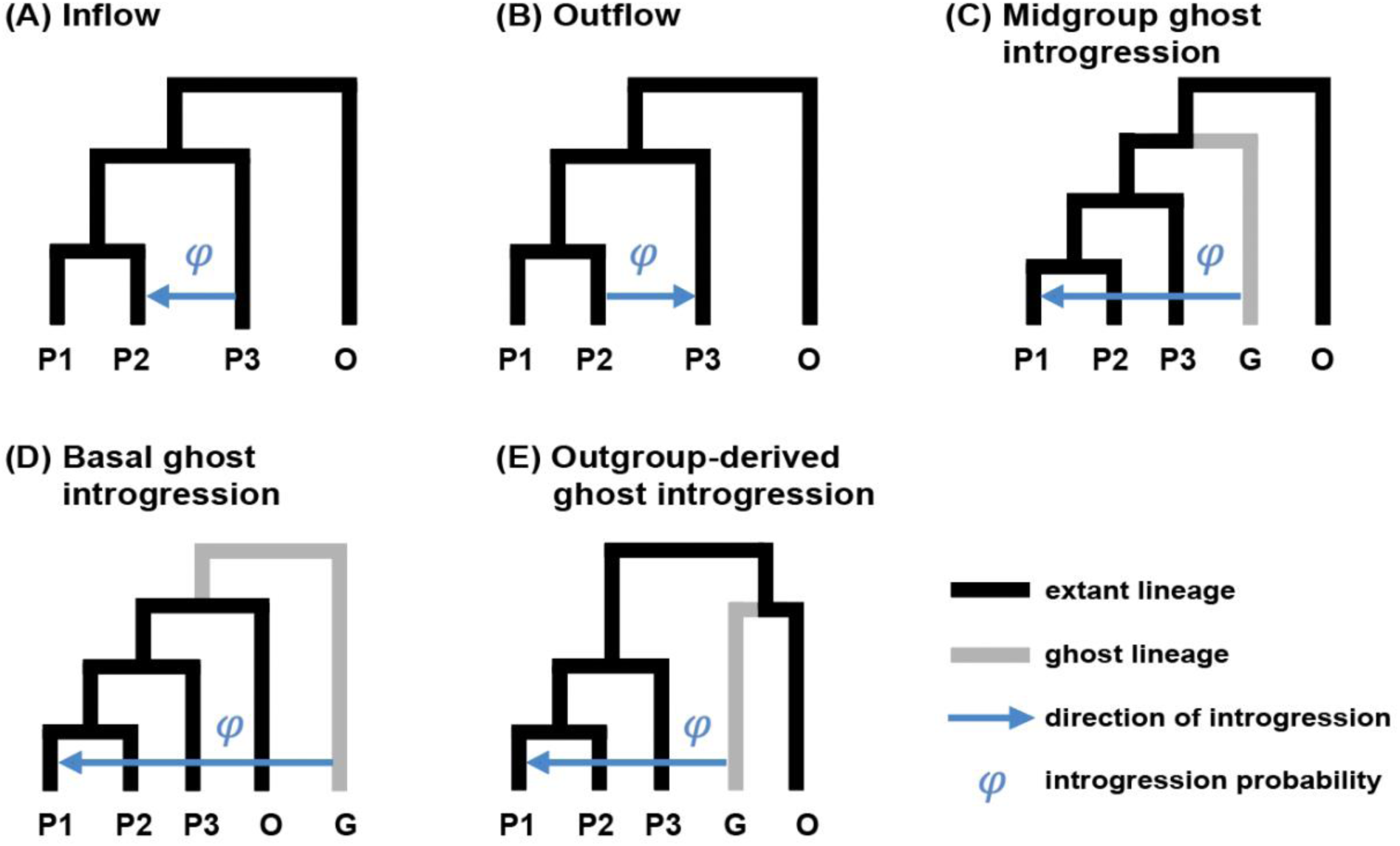
Five introgression scenarios that can produce a significantly positive D-statistic in a species quartet (((P1, P2), P3), O). (A, B) Introgression between non-sister taxa: gene flow from P3 into P2 (inflow, A) and from P2 into P3 (outflow, B). (C–E) Ghost introgression: the donor lineage may occupy one of three positions relative to the species tree—midgroup (C; between the ingroup clade and the outgroup), basal (D; diverging before the outgroup), or outgroup-derived (E; coalescing first with the outgroup). Each scenario generates an excess of ABBA over BABA site patterns, resulting in a significantly positive D-statistic.

In Step 3, each significant triple from Step 2 is sequentially evaluated using the Bayesian test (Ji et al. 2023) implemented in BPP (Flouri et al. 2020) to confirm genuine introgression events. The test evaluates support for gene flow (*H*₁) versus its absence (*H*₀) by computing a Bayes factor via the Savage–Dickey density ratio (Dickey 1971). Specifically, a “null interval” for the introgression probability is defined as *φ* < *ε* within the H₁ parameter space is used to represent *H*₀. The Bayes factor, is approximated as 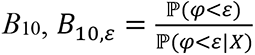, where the numerator and denominator represent prior and posterior probabilities of φ < ε, respectively; as *ε →* 0, this converges to the true Bayes factor *B*_10_. Following Ji et al. (Ji et al. 2023), we use *ε* = 0.01 and confirmed that *ε* = 0.001 yields similar results.

As outlined above, a single significant *D*-statistic can be explained by multiple alternative introgression events—namely, gene flow between non-sister ingroup taxa (inflow or outflow) or ghost introgression from an unsampled lineage. Thus, for each significant signal, we test three possible introgression events: (1) introgression from P2 to P3 (outflow), (2) introgression from P3 to P2 (inflow), and (3) introgression from a ghost lineage to P1 (Fig. 1). Notably, the inflow and outflow are modeled jointly by a bidirectional introgression (Yang and Flouri, 2022), under the simplifying assumption that, if both directions occurred, they did so simultaneously. Only those with decisive support *B*₁₀ > 100 (analogous to a *p*-value threshold of 1% in frequentist hypothesis testing) are retained.

Model construction proceeds iteratively. Starting with the highest-*D*ₚ triple, we incorporate the supported introgression events into the overall backbone tree. Horizontal reticulations (inflow and outflow) are placed on the branches of extant lineages, while ghost lineages are connected to the last common ancestor (LCA) of the involved taxa. Three exceptions warrant special handling: (1) When two triples share overlapping taxa, their introgression signals are jointly assessed and consolidated onto shared ancestral branches, following a parsimony-guided approach analogous to the *f*-branch method (Malinsky et al. 2021); (2) If multiple introgression events are inferred to fall on the same branch, their relative timing is determined by comparing the marginal likelihoods of the models with alternative orders (see step 4 below). (3) When the LCA is the ingroup root, all three positional scenarios of the ghost lineage relative to the outgroup (Fig. 2C–E) are also evaluated through marginal likelihood comparisons. Once a reticulation is incorporated for a given triple, any remaining triples whose signals are fully explained by existing events are excluded to avoid redundancy (Fig. 1); This process continues until all significant signals under the given species-tree topology are either incorporated or explained, yielding a finalized introgression model for that topology. The same steps are repeated for each starting tree, producing a set of introgression models corresponding to alternative species-tree topologies.

In Step 4, we compare the marginal likelihoods of introgression models constructed under each candidate species tree topology to identify the best-supported network. These marginal likelihoods are calculated in BPP via thermodynamic integration with Gaussian quadrature (Lartillot and Philippe 2006; Rannala and Yang 2017). The model with the highest marginal likelihood—representing the best-supported divergence and introgression scenario—is selected as the most probable under the MSci framework. Finally, we run the MCMC algorithm in BPP (Flouri et al. 2020) to obtain posterior parameter estimates for the best introgression model.

### Practical Considerations and Applications of D-BPP

A key consideration in implementing D-BPP is whether to include the outgroup that has been used in the *D*-statistic test in the BPP analyses. Incorporating an outgroup enables calibration of divergence times (e.g., the *Thuja* dataset in the below), which is particularly useful for fossil-calibrated studies, or for comparison with prior studies (the *Panthera* dataset; see Fig. S1). However, it also introduces uncertainty in ghost lineage placement, thereby increasing model complexity (Fig. 2).

As illustrated in Fig. 2, ghost lineages may occupy one of several positions relative to the outgroup: they may diverge between the ingroup and the designated outgroup (“midgroup”, Fig. 2C), before the outgroup divergence (“basal”, Fig. 2D), or may coalesce directly with the outgroup (“outgroup-derived”, Fig. 2E). Each of these configurations can produce a significant *D*-statistic (for mathematical proof see Supplementary Text). Misplacement of the ghost lineage can bias the inferred network topology and influence downstream parameter estimates. For instance, if a basal ghost lineage is incorrectly modeled as a midgroup donor, it may be anchored to the outgroup divergence time, leading to spurious timing and ancestry estimates.

Therefore, unless the outgroup is required for specific purposes (e.g., time calibration), we recommend excluding it from the BPP analysis within D-BPP to reduce computational burden and avoid biologically implausible ghost lineage placements under the MSci framework

Another practical consideration concerns how to incorporate multiple candidate introgression events into the model, either simultaneously or stepwise. When the number of significant *D*-statistics is small, it is feasible to adopt the strategy proposed by Ji et al. (Ji et al. 2025), in which all candidate events (3×*n*, where *n* is the number of species triples exhibiting significant *D*-statistic) are included in an initial full model, and unsupported events are subsequently pruned based on Bayes factor support. However, when *n* is large (say, *n* > 5), evaluating all possible combinations becomes computationally prohibitive. In such cases, we recommend the stepwise approach described by Ji et al. (Ji et al. 2023), in which introgression events are added sequentially in order of descending *D*_p_ values. As the number of parameters increases, so does the required MCMC chain length for reliable convergence, resulting in a substantial computational burden. The stepwise procedure mitigates this issue by introducing events sequentially, allowing for more efficient inference and better model tractability.

### The case of big cats (*Panthera*; Felidae)

We first analyzed the *Panthera* dataset from Santos et al. (2025), which includes 4,862 loci spanning five extant species—jaguar (*P. onca*), leopard (*P. pardus*), lion (*P. leo*), tiger (*P. tigris*), and snow leopard (*P. uncia*). Two individuals were sampled for jaguar, lion, and leopard, while tiger and snow leopard were each represented by a single genome. The domestic cat reference genome (v9) was used as the outgroup. Heterozygous sites were resolved using consensus base calling during mapping to the cat genome, resulting in pseudohaploid representations. Santos et al. (2025) considered three species tree topologies for jaguar, leopard, and lion: Topology 1 groups lion and leopard as sisters (Fig. 3A); Topology 2 places lion and jaguar as sisters (Fig. 3B); and Topology 3 unites leopard and jaguar (Fig. 3C).

**Fig. 3.**
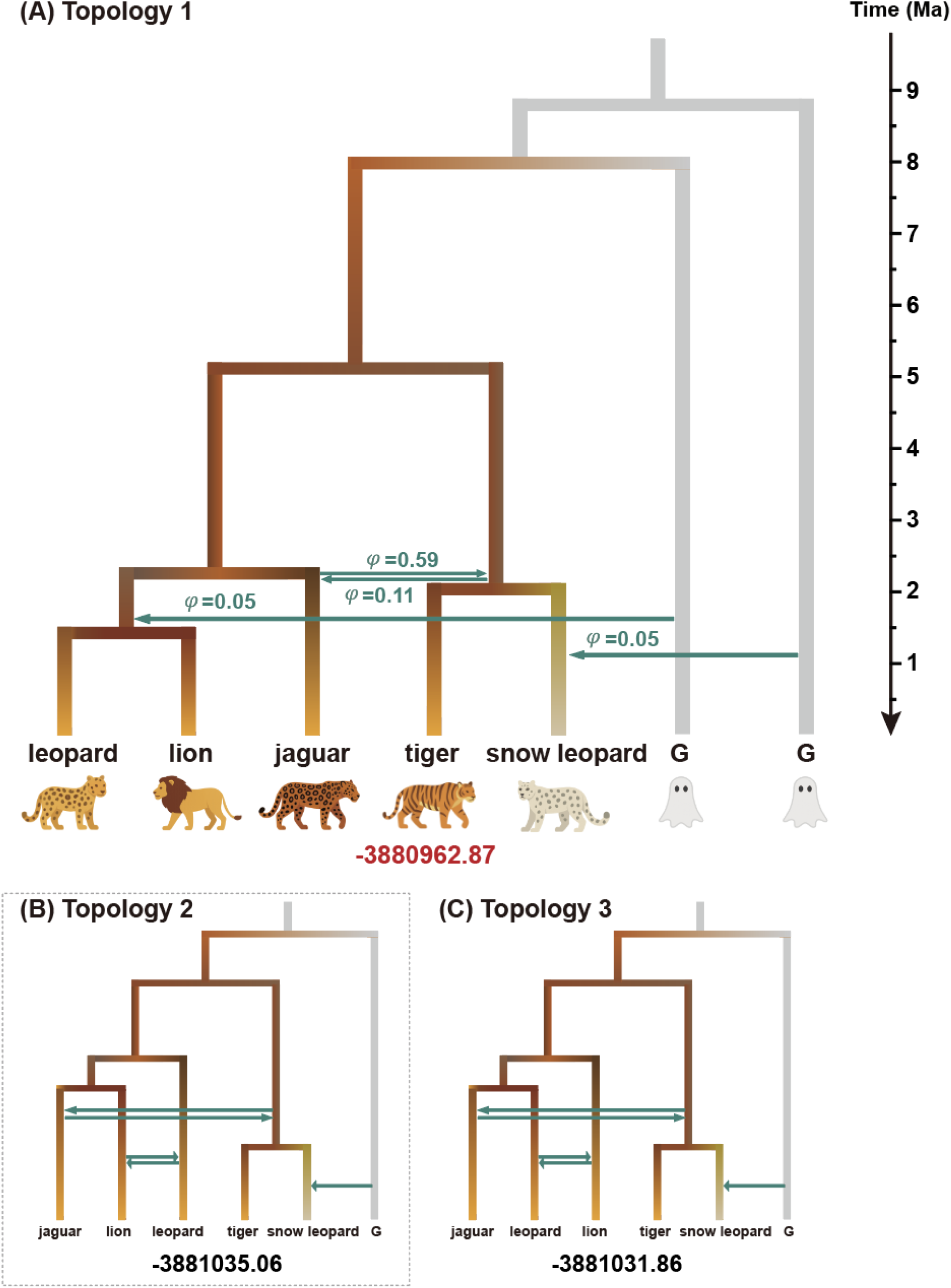
Introgression models under three alternative species-tree topologies for *Panthera*. (A) The introgression model under Topology 1, which also represents the best-supported model. The species tree is shown with yellow solid lines, while ghost lineages ("G") are indicated by the grey solid lines. Branch lengths are scaled to divergence time in millions of years (Ma), calibrated using a crown *Panthera* root age of 5.20 Ma (Santos et al. 2025). Introgression events are represented by dashed blue-green arrows and labeled with their corresponding introgression probabilities (*φ*). The marginal likelihood (log scale) of this model is indicated in red at the bottom. (B, C) Alternative introgression models inferred under topology 2 and topology 3, respectively.

For each topology, we constructed an introgression model. All 4,862 loci were concatenated and converted to VCF format using SNP-sites (Page et al. 2016) for *D*- statistic tests in Dsuite (Malinsky et al. 2021). Each topology yielded multiple significant signals (*p*-value < 0.01), indicative of gene flow (Table 1). Owing to computational limitations, we conducted BPP modeling on two subsets defined by Santos et al. (2025): 2,455 loci (primary dataset) and 2,407 loci (validation dataset). All analyses used the primary dataset, with consistency checks against the validation dataset.

**Table 1.** *D*-Statistic and *D_p_* Values for Introgression Detection Based on the three topologies of *Panthera*.

| Tree | P1 | P2 | P3 | $D$-statistic | Z-score | p-value | $D_p$ |
| --- | --- | --- | --- | --- | --- | --- | --- |
| Topology 1 | snow leopard | tiger | jaguar | 0.24 | 8.75 | 2.30E-16 | 0.120 |
|  | snow leopard | tiger | leopard | 0.19 | 6.28 | 3.48E-10 | 0.089 |
|  | lion | jaguar | tiger | 0.18 | 6.45 | 1.12E-10 | 0.074 |
|  | leopard | jaguar | tiger | 0.17 | 6.81 | 9.57E-12 | 0.070 |
|  | snow leopard | tiger | lion | 0.15 | 4.63 | 3.61E-06 | 0.068 |
|  | leopard | jaguar | snow leopard | 0.11 | 5.31 | 1.10E-07 | 0.039 |
|  | lion | jaguar | snow leopard | 0.05 | 2.87 | 4.07E-03 | 0.020 |
|  | leopard | lion | snow leopard | 0.07 | 2.32 | 0.02 | — |
|  | lion | leopard | tiger | 0.01 | 0.25 | 0.80 | — |
|  | leopard | lion | jaguar | 0.01 | 0.25 | 0.80 | — |
| Topology 2 | jaguar | lion | leopard | 0.37 | 13.19 | 2.30E-16 | 0.281 |
|  | snow leopard | tiger | jaguar | 0.24 | 8.75 | 2.30E-16 | 0.120 |
|  | snow leopard | tiger | leopard | 0.19 | 6.28 | 3.48E-10 | 0.089 |
|  | lion | jaguar | tiger | 0.18 | 6.45 | 1.12E-10 | 0.074 |
|  | leopard | jaguar | tiger | 0.17 | 6.81 | 9.57E-12 | 0.070 |
|  | snow leopard | tiger | lion | 0.15 | 4.63 | 3.61E-06 | 0.068 |
|  | leopard | jaguar | snow leopard | 0.11 | 5.31 | 1.10E-07 | 0.039 |
|  | lion | jaguar | snow leopard | 0.05 | 2.87 | 4.07E-03 | 0.020 |
|  | leopard | lion | snow leopard | 0.07 | 2.32 | 0.02 | — |
|  | lion | leopard | tiger | 0.01 | 0.25 | 0.80 | — |
| Topology 3 | jaguar | leopard | lion | 0.36 | 16.97 | 2.30E-16 | 0.277 |
|  | snow leopard | tiger | jaguar | 0.24 | 8.75 | 2.30E-16 | 0.120 |
|  | snow leopard | tiger | leopard | 0.19 | 6.28 | 3.48E-10 | 0.089 |
|  | lion | jaguar | tiger | 0.18 | 6.45 | 1.12E-10 | 0.074 |
|  | leopard | jaguar | tiger | 0.17 | 6.81 | 9.57E-12 | 0.070 |
|  | snow leopard | tiger | lion | 0.15 | 4.63 | 3.61E-06 | 0.068 |
|  | leopard | jaguar | snow leopard | 0.11 | 5.31 | 1.10E-07 | 0.039 |
|  | lion | jaguar | snow leopard | 0.05 | 2.87 | 4.07E-03 | 0.020 |
|  | leopard | lion | snow leopard | 0.07 | 2.32 | 0.02 | — |
|  | lion | leopard | tiger | 0.01 | 0.25 | 0.80 | — |

Under Topology 1, we identified seven significant species triples with *D*_p_ values ranging from 0.120 to 0.02 (Table 1). For subsequent BPP analyses, we excluded the outgroup (domestic cat) and used the primary dataset (2455 loci) of Santos et al. (2025). We followed their parameter settings: the Jukes-Cantor (JC) substitution model applied to all loci; inverse-gamma priors for population size parameters (θ ∼ *IG* (3, 0.003)) and the root divergence time (τ_₀_ ∼ *IG* (3, 0.024)); and a uniform prior *U* (0, 1) for the introgression probability (*φ*). Each MCMC run included a burn-in of 8×10⁴×*k* iterations, followed by 8×10⁴×*k* samples drawn every 10 iterations, where *k* is the number of introgression events in the MSci model.

Beginning with the highest-*D*_p_ triple ((snow leopard, tiger), jaguar), we tested three introgression scenarios: (1) outflow from tiger to jaguar, (2) inflow from jaguar to tiger, and (3) ghost introgression to snow leopard. Only the ghost introgression scenario received decisive support (*B*_10_ > 100). We removed triples ((snow leopard, tiger), leopard) and ((snow leopard, tiger), lion), as both were explainable by the ghost introgression into snow leopard. The remaining four significant triples consistently involved lion or leopard as P1, jaguar as P2, and tiger or snow leopard as P3 (Table 1). By applying parsimony, we assigned the ancestor of lion and leopard as P1, and the ancestor of tiger and snow leopard as P3. We tested: (1) outflow from jaguar to P3-ancestor, (2) inflow from P3-ancestor to jaguar, and (3) ghost introgression to P1- ancestor. All three received decisive support (*B*_10_ > 100). Thus, the final model incorporated 3 introgression events: (1) ghost introgression to snow leopard, (2) a bidirectional introgression between jaguar and the ancestor of tiger and snow leopard, and (3) ghost introgression to the ancestor of lion and leopard (Fig. 3A).

Following essentially the same procedure as described above for Topology 1, we obtain the corresponding introgression models for Topology 2 (Fig. 3B) and Topology 3 (Fig. 3C), with details being given in Supplementary Text.

To identify the most appropriate introgression model, we compared the marginal likelihoods of the three final models inferred under the three alternative topologies. Marginal likelihoods were estimated using thermodynamic integration with Gaussian quadrature (Lartillot and Philippe 2006; Rannala and Yang 2017), implemented with 16 quadrature points. The model under Topology 1 showed a markedly higher marginal likelihood (−3,880,962.87) compared to those of Topology 2 (−3,881,035.06) and Topology 3 (−3,881,031.86), strongly supporting Topology 1-based network as the best-fitting model (Fig. 3). The same conclusion was reached using the alternative dataset of 2,407 loci. Notably, the best-supported network derived from Topology 1 involves bidirectional introgression between the jaguar and the ancestor of the tiger and snow leopard. This model is theoretically indistinguishable from an alternative in which the positions of the jaguar and this ancestral lineage are exchanged (Yang and Flouri 2022). However, such a scenario was not evaluated, as leopard, lion, and jaguar were constrained to form a monophyletic clade in our analysis.

Based on the estimated crown age of *Panthera* at 5.2 million years ago (Ma) in Santos et al. (2025), we calibrated relative divergence times into absolute estimates. The divergence between the ghost lineages and the common ancestor of all extant *Panthera* species was estimated to have occurred at ∼8.86 Ma and ∼8.02 Ma, respectively (Fig. 3A, Table S1). The split within the clade consisting of jaguar, lion, and leopard was inferred to have occurred ∼2.34 Ma, while the divergence between tiger and snow leopard was estimated to occur at ∼3.12 Ma (Fig. 3A, Table S1). The bidirectional introgression between jaguar and the ancestral lineage of tiger and snow leopard was estimated to have occurred ∼ 2.32 million years ago. The introgression probability from the jaguar into the ancestor of tiger and snow leopard was 0.59, whereas the reverse introgression—from the ancestor into the jaguar—was 0.11. The ghost introgression into the ancestor of lion and leopard was estimated to have occurred at ∼1.59 Ma with an introgression probability of 0.05 (Fig. 3A, Table S1).

The divergence between lion and leopard was estimated at ∼1.51 Ma. The ghost introgression to snow leopard was estimated to have occurred at ∼1.11 Ma with an introgression probability of 0.05 (Fig. 3A, Table S1).

To assess the performance of D-BPP in detecting complex introgression events, we simulated multilocus sequence data under the best-supported introgression model for big cats (Fig. 3A). The simulated dataset comprised five ingroup species—A (leopard), B (lion), C (jaguar), D (tiger), and E (snow leopard)—and an outgroup (O). Each locus included two sequences for species A, B, and C, and one sequence for D, E, and O. Using the JC69 model (Jukes and Cantor 1969) implemented in BPP (Flouri et al. 2020), we generated 5,000 independent loci (1,000 bp each), matching the scale of our empirical dataset. Initial screening for introgression signals was performed using *D*-statistics (Supplementary Text, Table S2). For Bayesian model validation and introgression inference in BPP, we analyzed a subset of 2,500 loci (outgroup excluded), following the same approach as in the above empirical analyses. D-BPP successfully recovered all four simulated introgression events (Supplementary Text, Table S3), with reticulation nodes accurately placed on the corresponding species tree branches. These results demonstrate D-BPP’s robustness in detecting and characterizing hybridization events while resolving complex evolutionary histories in big cats.

### The case of arborvitae (*Thuja*; Cupressaceae)

We next analyzed the *Thuja* dataset from Li et al. (2022), which comprises 1,196 loci from 20 individuals, with *Thujopsis dolabrata* (2 individuals) as the outgroup. The dataset included *T. plicata* (4), *T. occidentalis* (4), *T. koraiensis* (4), *T. sutchuenensis* (3), and *T. standishii* (3). In their study, Li et al. (2022) consistently grouped *T. sutchuenensis* and *T. standishii* as sister species, while *T. plicata*, *T. occidentalis* and *T. koraiensis* formed a monophyletic clade. However, this clade exhibited extensive gene tree discordance (Li et al. (2022): Fig. 1a), indicating considerable phylogenetic uncertainty. To account for this, we evaluated three alternative species tree topologies involving *T. plicata*, *T. occidentalis* and *T. koraiensis*, while consistently treating *T. sutchuenensis* and *T. standishii as* a sister clade to the three species (Fig. 4). Note that Topology 1 is the inferred species tree under the assumption of no gene flow (Li et al. 2022).

**Fig. 4.**
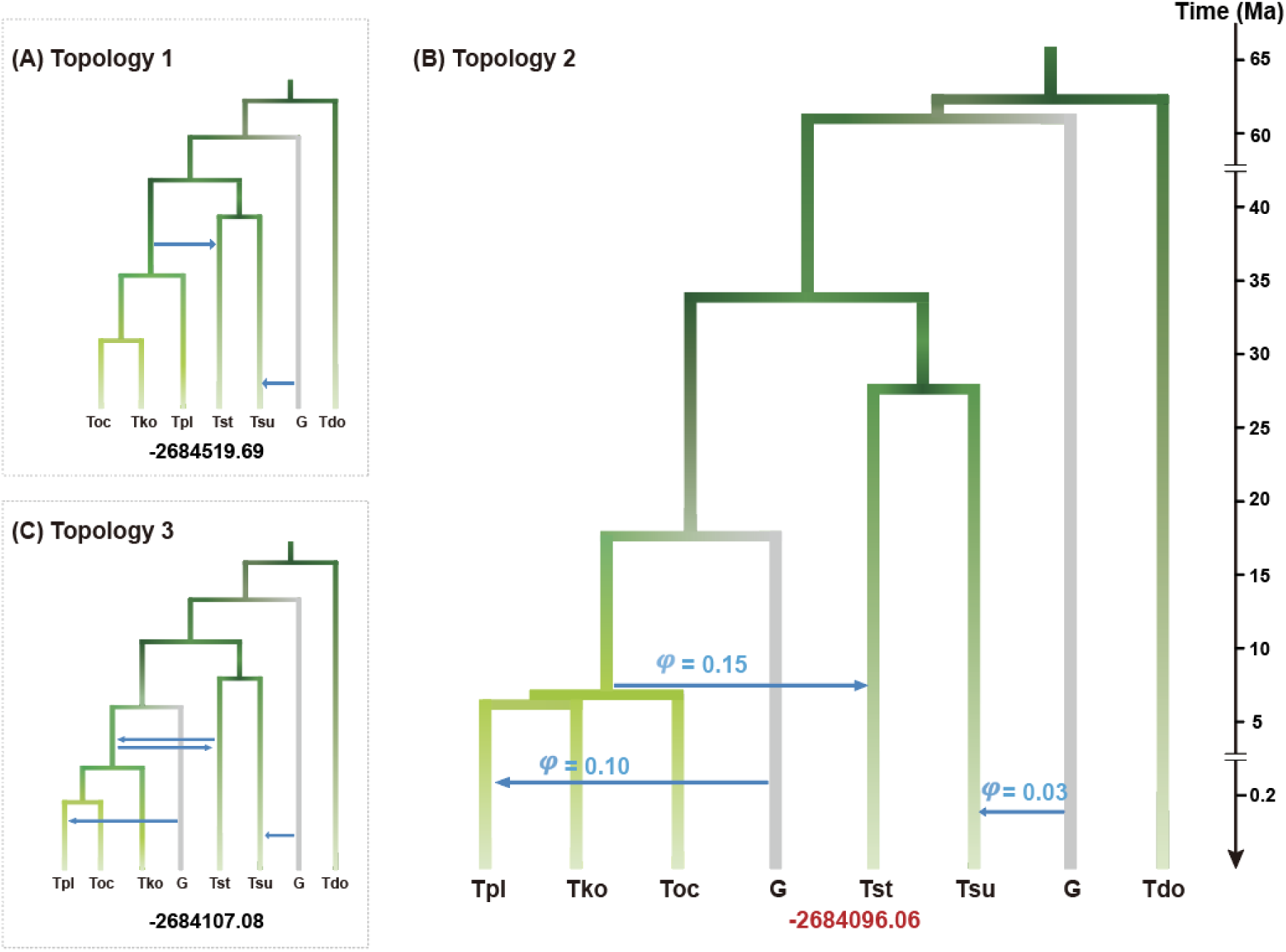
Introgression models under three alternative species-tree topologies for *Thuja*. (A, C) Introgression models inferred under topology 1 and topology 3, respectively, with supported introgression events and their corresponding marginal likelihoods. (B) Introgression model under topology 2, identified as the best-supported model. Green solid lines represent the species tree, while grey lines denote ghost lineages (“G”). All branch lengths are scaled to absolute divergence times (in millions of years, Ma), calibrated using a root age of 62.68 Ma (Li et al. 2022). Introgression events are indicated by dashed blue arrows and annotated with their estimated introgression probabilities (*φ*). The marginal likelihood (log scale) of this model is shown in red at the bottom.

D-statistic tests identified at least four significant introgression signals (*p*-value < 0.01) for each species-tree topology, supporting a complex reticulated evolutionary history (Table 2). We evaluated introgression under the three alternative species trees using the D-BPP framework (see Supplementary Text for details). Following Li et al. (2022), we modeled the ghost lineage as a midgroup donor and retained the outgroup (*Thujopsis dolabrata*) in our BPP analyses to allow calibration of divergence times.

**Table 2.** *D*-Statistic and *D_p_* Values for Introgression Detection Based on the three topologies of *Thuja*.

| Tree | P1 | P2 | P3 | *D*-statistic | Z-score | p-value | *D<sub>p</sub>* |
| --- | --- | --- | --- | --- | --- | --- | --- |
| Topology 1 | Tsu | Tst | Tko | 0.42 | 9.79 | 2.30E-16 | 0.246 |
|  | Tsu | Tst | Tpl | 0.40 | 6.85 | 7.15E-12 | 0.229 |
|  | Tsu | Tst | Toc | 0.40 | 7.09 | 1.35E-12 | 0.227 |
|  | Toc | Tko | Tsu | 0.10 | 3.17 | 1.51E-03 | 0.039 |
|  | Toc | Tko | Tst | 0.09 | 2.17 | 0.03 | — |
|  | Tpl | Tko | Tst | 0.07 | 1.39 | 0.17 | — |
|  | Tpl | Tko | Tsu | 0.05 | 0.80 | 0.42 | — |
|  | Toc | Tpl | Tsu | 0.05 | 0.86 | 0.39 | — |
|  | Toc | Tko | Tpl | 0.02 | 0.33 | 0.74 | — |
|  | Toc | Tpl | Tst | 0.01 | 0.16 | 0.87 | — |
| Topology 2 | Tsu | Tst | Tko | 0.42 | 9.79 | 2.30E-16 | 0.246 |
|  | Tsu | Tst | Tpl | 0.40 | 6.85 | 7.15E-12 | 0.229 |
|  | Tsu | Tst | Toc | 0.40 | 7.09 | 1.35E-12 | 0.227 |
|  | Tpl | Tko | Toc | 0.18 | 4.01 | 6.14E-05 | 0.130 |
|  | Toc | Tko | Tsu | 0.10 | 3.17 | 1.51E-03 | 0.039 |
|  | Toc | Tko | Tst | 0.09 | 2.17 | 0.03 | — |
|  | Tpl | Tko | Tst | 0.07 | 1.39 | 0.17 | — |
|  | Tpl | Tko | Tsu | 0.05 | 0.80 | 0.42 | — |
|  | Toc | Tpl | Tsu | 0.05 | 0.86 | 0.39 | — |
|  | Toc | Tpl | Tst | 0.01 | 0.16 | 0.87 | — |
| Topology 3 | Tsu | Tst | Tko | 0.42 | 9.79 | 2.30E-16 | 0.246 |
|  | Tsu | Tst | Tpl | 0.40 | 6.85 | 7.15E-12 | 0.229 |
|  | Tsu | Tst | Toc | 0.40 | 7.09 | 1.35E-12 | 0.227 |
|  | Tpl | Toc | Tko | 0.17 | 4.64 | 3.55E-06 | 0.121 |
|  | Toc | Tko | Tsu | 0.10 | 3.17 | 1.51E-03 | 0.039 |
|  | Toc | Tko | Tst | 0.09 | 2.17 | 0.03 | — |
|  | Tpl | Tko | Tst | 0.07 | 1.39 | 0.17 | — |
|  | Tpl | Tko | Tsu | 0.05 | 0.80 | 0.42 | — |
|  | Toc | Tpl | Tsu | 0.05 | 0.86 | 0.39 | — |
|  | Toc | Tpl | Tst | 0.01 | 0.16 | 0.87 | — |

The final model under Topology 1 included two gene-flow events: (1) introgression from the ancestor of *T. koraiensis*, *T. plicata*, and *T. occidentalis* to *T. standishii*; and (2) ghost introgression to *T. sutchuenensis* (Fig. 4A). Under Topology 2, three events were supported: (1) ghost introgression into *T. sutchuenensis*; (2) introgression from the same ancestor of *T. koraiensis*, *T. plicata*, and *T. occidentalis* to *T. standishii*; and (3) ghost introgression into *T. plicata* (Fig. 4B). Topology 3 yielded four events: (1) ghost introgression into *T. sutchuenensis*; (2) introgression from the ancestor of *T. koraiensis*, *T. plicata*, and *T. occidentalis* to *T. standishii*; (3) introgression from *T. standishii* to this ancestor; and (4) ghost introgression into *T. plicata* (Fig. 4C).

To identify the most appropriate introgression model, we compared the marginal likelihoods of the three introgression models inferred under the three topologies. The introgression model of Topology 2 exhibited a substantially higher marginal likelihood (−2,684,096.06) compared to that of Topology 1 (−2,684,519.69) and Topology 3 (−2,684,107.08), strongly supporting Topology 2 as the best species tree for modeling introgression (Fig. 4).

We calibrated the root age (divergence between *Thuja* and the outgroup *Thujopsis dolabrata*) at 62.68 Ma as suggested in Li et al. (2022). The divergence between the ghost lineage introgressing into *T. sutchuenensis* and the common ancestor of all extant *Thuja* species was estimated to have occurred at ∼61.37 Ma, while the stem lineage leading to extant *Thuja* species diverged at ∼34.18 Ma (Fig. 4B, Table S4). The divergence time between *T. sutchuenensis* and *T. standishii* was estimated at ∼27.94 Ma and the ghost lineage introgressing into *T. plicata* diverged in ∼17.96 Ma (Fig. 4B, Table S4). The time of introgression from the ancestor of *T. occidentalis*, *T. koraiensis*, and *T. plicata* to *T. standishii* was estimated at ∼6.91 Ma with an introgression probability of 0.15. The divergence among these three species (*T. occidentalis*, *T. koraiensis*, and *T. plicata*) was inferred to have occurred essentially at the same time (∼6.91 Ma), i.e., nearly forming a polytomy. The ghost introgression into *T. sutchuenensis* was estimated to have occurred at ∼ 0.24 Ma with a small introgression probability of 0.03, and that into *T. plicata* at ∼0.16 Ma with a moderate introgression probability of 0.10 (Fig. 4B, Table S4).

## Discussion

In this study, we developed and implemented D-BPP, a unified framework that integrates the computational efficiency of the heuristic *D*-statistic with the statistical efficiency of full-likelihood Bayesian inference (BPP) to accurately infer complex phylogenetic networks involving gene flow from both extant and ghost lineages.

While traditional *D*-statistic method is effective in determining the presence or absence of introgression events, it cannot distinguish between scenarios such as ghost introgression or gene flow between non-sister taxa (Tricou et al. 2022b; Pang and Zhang 2024). In contrast, the full-likelihood Bayesian test in BPP offers high power for identifying specific introgression scenarios, despite at relatively substantial computational cost (Ji et al. 2023). By combining heuristic and Bayesian methods into a single framework for tree-based inference of phylogenetic networks, D-BPP efficiently searches for candidate introgression events and rigorously validates them to construct a well-supported introgression model. When applied to simulated multilocus sequence data generated from the inferred reticulate evolutionary history of big cats, D-BPP recovered all introgression events, indicating its potential as an effective tool for inferring complex phylogenetic networks. Additionally, by screening alternative species-tree topologies through Bayesian model comparison of their corresponding introgression models, D-BPP enables the joint inference of species divergence history and gene flow patterns.

### Methodological innovations behind D-BPP

D-BPP introduces several key methodological innovations. First, it offers a statistically robust yet computationally tractable framework for resolving complex networks without requiring exhaustive searches across model space. Existing heuristic methods, such as—SNaQ (Solís-Lemus and Ané 2016), NANUQ (Allman et al. 2019), and PhyNEST (Kong et al. 2025)—accelerate computation via pseudo-likelihood or distance-based approaches but are generally restricted to level-1 networks. As a result, they tend to produce oversimplified topologies that overlook hybridization signals detectable by *D*-statistics. In contrast, D-BPP can infer complex level-*k* (*k* > 1) networks, thereby achieving greater consistency between inferred topologies and genome-wide introgression signals. Our empirical analyses of the *Panthera* and *Thuja* datasets demonstrate that D-BPP captures higher-level network complexity, including prevalent ghost introgression events (Figs. 3, 4), which are notoriously difficult to recover without leveraging full-likelihood approaches.

Second, D-BPP addresses ghost introgression—an often-overlooked but biologically significant aspect of reticulate evolution—by explicitly incorporating it into model inference. Previous studies of *Panthera* (Figueiró et al. 2017; Santos et al. 2025) and *Thuja* (Li et al. 2022), which relied on *D*-statistics or its variants such as *D*_FOIL_ (Pease and Hahn 2015), as well as heuristic network methods, either entirely ignored ghost lineages or captured only a limited subset of such events, likely due to constraints in model identifiability (Pang and Zhang 2024). By contrast, the D-BPP framework overcomes this limitation, enabling quantification of ghost introgression prevalence in nature while avoiding spurious inferences of horizontal gene flow between non-sister ingroups—a known artifact of ignoring ghost lineages (Tricou et al. 2022b).

Crucially, while introgression is more likely among closely related species, hybridization and introgression are also well documented between highly divergent species in plants (e.g., (Stull et al. 2020)) and animals (e.g., (Tea et al. 2020)).

Therefore, phylogenetically distant and unsampled donors—i.e., ghost lineages—may be an inevitable source of introgressed alleles in many empirical systems.

Finally, a key strength of D-BPP lies in its ability to accurately recover species divergence histories despite pervasive gene flow. Empirical phylogenomic studies often struggle to infer reliable species trees under widespread ILS and introgression (Jiao et al. 2020; Pang and Zhang 2023), sometimes producing conflicting topologies that complicate introgression inference. D-BPP addresses this challenge by evaluating competing species-tree topologies and selecting the one associated with the best-supported introgression model, under the assumption that it most accurately reflects both speciation and reticulation history. Using this strategy, D-BPP recovered the lion-leopard sister relationship in *Panthera* (contrasting with Santos et al. (2025)) and the *T. occidentalis–T. koraiensis* sister relationship in *Thuja* (differing from Li et al. (2022)), demonstrating its potential for joint inference of speciation and introgression histories.

### Insights into the species tree estimation and reticulate evolution in *Panthera* and *Thuja*

The application of the D-BPP framework to the *Panthera* and *Thuja* datasets showcases its utility in reconstructing complex species relationships and reticulate evolutionary histories. In the big cat genus *Panthera*, our results support an introgression model that retains the traditional species tree topology—grouping lion and leopard as sister taxa. The model incorporates two ghost introgression events and a bidirectional gene flow between jaguar and the ancestor of tiger and snow leopard (Fig. 3A). This finding stands in sharp contrast to Santos et al. (2025), who rejected the lion–leopard sister relationship as the true species topology, attributing its frequent recovery along chromosomes to extensive post-speciation gene flow.

However, their analysis was limited to a subset of taxa, focusing on introgression only among lion, leopard, and jaguar, without fully considering introgression involving the other two *Panthera* species (i.e., tiger and snow leopard) or ghost lineages. In contrast, our study employed the D-BPP framework to jointly analyze all five *Panthera* species, considering introgressions from both extant and ghost lineages under three alternative species tree topologies. To ensure comparability with the network inference of Santos et al. (2025), we also included the outgroup (domestic cat) in our search for the best-supported introgression model. This model (Fig. S1) achieved a substantially higher marginal likelihood—exceeding the top introgression model from Santos et al. (2025) by over 150 log units—indicating a much better fit to the data. Importantly, these results demonstrate that the lion-leopard sister relationship is statistically the best-supported species-tree topology once comprehensive introgression scenarios are considered—i.e., as it incorporates ghost introgression and expands beyond the limited taxon subset considered previously.

We report, for the first time, evidence of ghost introgression to the snow leopard (*Panthera uncia*) (Fig. 3A). The phylogenetic position of the snow leopard has long been debated due to its distinctive morphology and adaptations to high-altitude environments. Initially classified as the sole member of *Uncia* (Christiansen 2008), molecular phylogenies later reclassified it within *Panthera* as sister to tiger (*P. tigris*) (Johnson et al. 2006; Li et al. 2016; Figueiró et al. 2017). However, persistent discordance between molecular and morphological data has remained unresolved. The ghost introgression inferred here offers a plausible explanation for this discrepancy, potentially reflecting gene flow from an unsampled donor lineage that introduced unique genomic signatures into the snow leopard. While further research is needed, these findings illustrate how ghost introgression can help reconcile longstanding inconsistencies in the evolutionary history of snow leopard.

In *Thuja*, our D-BPP analysis revealed a species divergence history and reticulate evolutionary pattern distinct from those reported by Li et al. (2022) (Fig. 4). We recovered *T. koraiensis*, *T. plicata* and *T. koraiensis* effectively as a polytomy (Fig. 4B), contrasting with Li et al. (2022) ’s treatment of *T. koraiensis* and *T. occidentalis* as sister lineages. This topological discrepancy likely stems from the combined effects of strong incomplete lineage sorting (ILS)—evidenced by the near-simultaneous divergence among *T. occidentalis*, *T. koraiensis*, and *T. plicata* in our network—and ghost introgression into *T. plicata*. These results are consistent with theoretical predictions by Pang and Zhang (2023), who showed that even low levels of ghost introgression can distort species tree topology under conditions of strong ILS. Beyond correcting the topology, our analysis identified an additional ghost introgression event and a horizontal gene flow event missed by Li et al. (2022), who reported only one ghost introgression (to *T. sutchuenensis*). Together, these results suggest a history of persistent gene exchange in *Thuja*, shaped by ancient hybridization and lineage extinction—patterns that align with its East Asia–North America disjunction and status as a Tertiary relict. Given the extensive range shifts, climatic fluctuations, and extinction events associated with such lineages, ghost introgression is particularly plausible in relict plant clades (Milne and Abbott 2002; Milne 2006).

The detection of ghost introgression in *Panthera* and *Thuja* highlights its prevalence across diverse taxa. As emphasized by (Tricou et al. 2022a, b), its apparent rarity likely reflects methodological limitations rather than biological absence. Crucially, detectability depends only on introgression from an unsampled lineage (extinct or extant) into a sampled ingroup—irrespective of outgroup selection. This contrasts with Tricou et al. (2022b), who focused solely on midgroup ghosts. As shown in Fig. 2, ghost lineages can occupy three phylogenetic positions relative to the outgroup (midgroup, basal, or outgroup-derived), with the chosen outgroup’s position influencing which ghost types are more easily detected. When all ghost types are considered, presumably a greater proportion of significant *D*-statistics is attributable to ghost introgression than envisaged by Tricou et al. (2022b). Given that most empirical systems permit gene inflow from unknown sources, ghost introgression will usually be detected whenever sought. Aligning with this, recent studies report widespread ghost introgression in birds (Zhang et al. 2019), mammals (Kuhlwilm et al. 2019; Wang et al. 2020; Pawar et al. 2023), frogs (Shen et al. 2025), fishes (Baraf et al. 2025), plants (Luo et al. 2017; Ru et al. 2018; Ding et al. 2022; Li et al. 2022; Tiley et al. 2023; Zhang et al. 2024), and hominids (Green et al. 2010; Sankararaman et al. 2014; Rochaix 2022). These findings collectively underscore the need to explicitly account for ghost lineages in phylogenomic analyses, especially in ancient or extinction-prone clades.

## Limitations and Future Directions

Despite its strengths, D-BPP has several notable limitations. First, the range of detectable gene flow events is restricted to those capable of producing a significant *D*- statistic. As a result, the current implementation does not accommodate introgression between sister taxa, nor does it detect ghost introgression into P3 (rather than ghost→P1 shown in Fig. 2C-E). These omissions may lead to missed but biologically plausible scenarios. However, recent methodological developments in BPP (Ji et al. 2023) offer potential avenues to address these limitations in future implementations. Second, the requirement for stepwise manual intervention in searching for candidate introgression events may introduce investigator bias. Third, while computationally tractable for small to moderate datasets, the method’s computational demands scale nonlinearly with taxon number, making analyses involving more than 10 species particularly challenging.

A promising strategy to scale D-BPP for phylogenomic datasets involves adopting divide-and-conquer approaches (Zhu et al. 2019). By applying D-BPP to a simplified "backbone" dataset of representative taxa from major clades, model complexity can be reduced while retaining key evolutionary signals. This enables D-BPP to infer deep reticulation events without being confounded by within-clade noise. Focusing on exemplar taxa allows the method to efficiently scale to large phylogenies while maintaining interpretability of deep hybridization patterns. Such an exemplar-based approach would enhance the scalability of D-BPP for extensive taxon sets and clarify complex reticulate histories. However, because taxon subsampling increases the risk of introgression donors or recipients being unsampled, accounting for ghost introgression becomes even more imperative.

## Concluding Remarks

D-BPP offers a statistically rigorous yet computationally efficient framework for inferring phylogenetic networks under complex hybridization and introgression. By integrating the speed of heuristic *D*-statistics with the statistical rigor of full-likelihood Bayesian inference, D-BPP addresses key limitations of existing methods, producing more accurate reconstructions of species relationships and gene flow histories. Its consistent performance across different empirical datasets—including animals (*Panthera*) and plants (*Thuja*)—demonstrates its versatility and positions it as a premier tool for phylogenomic studies of reticulate evolution. Furthermore, by expanding the conceptual framework of ghost introgression beyond the midgroup scenario emphasized by Tricou et al. (2022b), D-BPP further reveals this phenomenon to be substantially more prevalent than previously recognized. Together, these advances position D-BPP as a significant methodological contribution, enhancing both empirical inference and theoretical understanding of reticulate evolution.

## Acknowledgments

We are grateful to the authors of *Panthera* (Santos et al. 2025) and *Thuja* (Li et al. 2022) for sharing the genomic sequence data with us. We thank Ziheng Yang, Yuttapong Thawornwattana, and Shou-Hsien Li for their constructive comments and suggestions that greatly improve the manuscript.

## Author Contributions

D.-Y.Z. and W.-N.B. conceived and designed the project; Y.Y., X.-X.P. and D.-Y.Z. developed the conceptual framework and wrote the paper; Y.Y. analyzed the data; W.-N.B., D.-Y.M., and B.-W.Z. revised and proofed the paper; All authors approved the final version.

## Data and materials availability

The datasets analyzed in this study are publicly available and were obtained from *Panthera* (Santos et al. 2025) and *Thuja* (Li et al. 2022).

